# A middle ground where executive control meets semantics: The neural substrates of semantic-control are topographically sandwiched between the multiple-demand and default-mode systems

**DOI:** 10.1101/2021.11.26.470178

**Authors:** Rocco Chiou, Elizabeth Jefferies, John Duncan, Gina F. Humphreys, Matthew A. Lambon Ralph

## Abstract

Semantic control is the capability to operate on meaningful representations, selectively focusing on certain aspects of meaning while purposefully ignoring other aspects based on one’s behavioural aim. This ability is especially vital for comprehending figurative/ambiguous language. It remains unclear, at the topographical level, why/how regions involved in semantic control seem reliably juxtaposed alongside other functionally specialised regions in the association cortex. We investigated this issue by characterising how semantic control regions topographically relate to the default-mode network (associated with memory and abstract cognition) and multiple-demand network (associated with executive control). Topographically, we confirmed that semantic control areas were sandwiched by the default-mode and multi-demand networks, forming an orderly arrangement observed both at the individual- and group-level. Functionally, semantic control regions exhibited ‘hybrid’ responses, fusing a preference for cognitively demanding processing (multiple-demand) and a preference for meaningful representations (default-mode) into a domain-specific preference for difficult operations on meaningful representations. When projected onto the principal gradient of human connectome, the neural activity of semantic control showed a robustly dissociable trajectory from visuospatial control, implying different roles in the functional transition from sensation to cognition. We discuss why the hybrid functional profile of semantic control regions might result from their intermediate topographical positions.

## Introduction

The human brain implements a wide variety of executive mechanisms to control behaviour flexibly; selectively deploying attention, suppressing unwanted/habitual reactions, actively maintaining contents in mind for later use (working memory), and swiftly adjusting when an error is detected. While different types of executive control processes differ greatly in how they are operated at the cognitive level (e.g., actively rehearsing number strings in memory entails disparate processes from attentively searching for an item in a messy space), they all engage the multiple-demand (MD) system (Duncan 2010; Duncan *et al*. 2020) – a set of frontoparietal regions that show heightened activation when attaining a behavioural goal becomes more difficult. There is robust evidence showing that neural activity of the MD system is sensitive to difficulty under miscellaneous tasks, stimuli types, and from different sensory modalities, supporting its ‘multiple-demand’ nature. In the present study, we investigated two issues of high relevance to the research disciplines of language, cognitive control, and connectomics: (***i***) We compared the topographical distribution of neural activities triggered by controlling semantic information vs. those triggered by controlling visuospatial information, examining whether semantic control similarly entails the MD system (like visuospatial control) or whether it requires additional regions (cf. Whitney *et al*. 2011; Gao *et al*. 2021). (***ii***) In the context of macroscale cortical organisation (cf. Margulies *et al*. 2016), we asked how regions involved in semantic control are spatially related to other well-studied cortical networks (the MD system and the default-mode network), since previous studies have suggested that semantic control regions might lie at the intersection of these networks and show intermediate responses and connectivity patterns (Davey et al., 2016; Wang *et al*., 2020). By investigating the topographical layout, we seek to clarify what inferences can be made regarding how semantic control regions acquire a ‘partially semantic, partially executive’ proclivity as a result of their unique topographical position.

We start by reviewing previous literature regarding the neural correlates of semantic control. Next, we review the recent development of human connectome research. Specifically, we discuss how a dimensionality-reduced framework of cortical gradients may provide useful explanation as to why the topography of semantic control are spatially configured the way they are, and how their locations may give rise to unique functions. Finally, we introduce the current experimental design and explain the logic of our analytical approaches.

### Prior literature regarding the neural mechanisms of semantic control

Operationally speaking, ‘semantic control’ can be defined as the ability to selectively access and manipulate meaningful information based on contextual demands – such as selectively interpreting *crane* as a bird rather than a type of machine (cf. Thompson-Schill *et al*. 1997; Badre *et al*. 2005).

When studying semantic control, researchers usually manipulate the difficulty of retrieving information, such as asking participants to access an infrequent meaning. This contrast of harder versus easier tasks mirrors the manipulation of difficulty across different domains used to probe the MD system (see Fedorenko *et al*., 2013). Many years of research has revealed that semantic control relies on a set of widely distributed regions (for meta-analysis, see Jackson 2020). Among these areas, the inferior frontal gyrus (IFG) and posterior middle temporal gyrus (pMTG) of the left hemisphere and the dorsomedial prefrontal cortex (dmPFC) have received most attention in previous research (e.g., Whitney *et al*. 2011; Noonan *et al*. 2013; Davey *et al*. 2016; Chiou *et al*. 2018; Wang *et al*. 2018; Gao *et al*. 2021). Interestingly, however, while some regions within the MD network are involved in semantic control, there is only *partial* overlap between the semantic control network and frontoparietal MD regions, and the majority of semantic control areas fall *outside* the MD system (Jackson 2020). This suggests that the MD system might not be the dominant contributing force to semantic control. There have also been many investigations regarding the role of MD regions in language-related tasks (for discussion, see Fedorenko and Shain 2021). While some MD regions become more active when the difficulty of accessing semantics was deliberately manipulated (Chiou *et al*. 2018; Gao *et al*. 2021), most regions of the MD network are minimally involved in *naturalistic* language comprehension, such as passively comprehending text/speech when there is no additional task requirement (e.g., Blank and Fedorenko 2017; Mineroff *et al*. 2018; Diachek *et al*. 2020; Wehbe *et al*. 2021; Malik-Moraleda *et al*. 2022). By contrast, semantic control areas (the LIFG and pMTG) are engaged under naturalistically comprehending circumstances. Interestingly, by visual inspection of activation maps, semantic control areas seem to lie adjacent to MD areas, often abutting each other, which forms a specific cortical tapestry. Here we aim to characterise this cortex distribution.

### Condensing the human-connectome into cortical gradients

Recent developments in brain cartography have demonstrated the utility of dimensionality-reduction methods that describe the whole-brain’s connectome in terms of a set of components (e.g., Margulies *et al*. 2016; Oligschläger *et al*. 2017; Bajada *et al*. 2019). Each component is called a ‘gradient’ as it characterises how cortical areas differ from one another along a graded dimension. The principal gradient, which explains most variance in connectivity, describes the gradual transition from regions involved in unimodal processes (e.g., the primary auditory, visual, motor cortex) to heteromodal cortex (e.g., the default-mode network). Margulies *et al*. (2016) projected the seven canonical resting-state networks (identified by Yeo *et al*. 2011) and NeuroSynth meta-analysis results onto the principal gradient and found an orderly arrangement – the unimodal end of the gradient supports sensory functions, while regions at the heteromodal end are involved in more abstract processes (such as memory or social cognition, which robustly activate the default-mode network). A key aim of our study is to establish where semantic control is situated on the principal gradient. Inspired by previous gradient analysis (cf. Murphy *et al*. 2018; Murphy *et al*. 2019; Wang *et al*. 2020), which have shown systematic functional changes along the gradient for memory-related processes, we projected the neural correlates of semantic control onto the gradient, and compared its location with the MD and default-mode networks. Previously, Margulies *et al*. (2016) found that MD areas (and studies focused on executive control) fall mostly in the middle of the principal gradient while semantic/language areas are located near the principal gradient’s heteromodal end. Here, we directly test the hypothesis that the locus of semantic control regions would be intermediate between the MD system and memory-related regions of the DMN.

### Experimental design and analytical approaches

We used task-based functional MRI and orthogonally manipulated two factors: the type of operation required by the task (Semantic *vs*. Visuospatial) and task difficulty (Easy *vs*. Hard). This 2-by-2 factorial design allowed us to identify three sets of brain regions (cf. Humphreys and Lambon Ralph 2017, for a similar approach): (**1**) areas showing the typical cortical morphology of the MD system, which favoured hard over easy operations in the visuospatial domain (see Mineroff et al. 2018); (**2**) areas of the default-mode system, which preferred easy to hard operations in the visuospatial domain; areas showing a ‘hybrid’ response profile: stronger activation for both *hard semantic* decisions and *easy visuospatial* decisions. This response suggests a fusion of semantic processing with executive control – i.e., to activate these regions, the task needed to be *both* semantic and difficult. We projected these areas to the principal gradient (Margulies *et al*., 2016) and compared the trajectory of semantic control along the gradient with the multiple-demand network and default-mode network.

To foreshadow the main findings, the multiple-demand and default-mode networks were situated at distinct loci on the principal gradient. Critically, semantic control areas were both functionally and topographically distinct from both the MD network and default-mode network. Topographically, semantic control regions occupied the intermediate territory between MD and default-mode regions on the cortical surface. Functionally, semantic control areas exhibited an interaction effect (showing responses to both semantic difficulty and visuospatial easiness). We suggest this intermediate location of the semantic control network allows executive control to coalesce with semantic memory (see Wang *et al*. 2021 for similar interpretations). This lends further support to the proposal that there is specialised machinery for controlling the retrieval of semantic information, dissociable from the generality of MD network (e.g., Davey *et al*. 2016; Chiou *et al*. 2018; Jackson 2020; Gao *et al*. 2021).

## Methods

### Participants

Twenty-five volunteers gave informed consent before the fMRI experiment. The sample consisted of 15 females and 10 males, with an average age = 28 years-old and SD = 6. All volunteers speak English as their mother tongue and are right-handed. All participants completed a screening questionnaire about magnetic resonance imaging safety before the scanning session began. None of them reported having any previous brain injury, neurological, or psychiatric condition. This study was reviewed and approved by the ethics board at University of Cambridge.

### Design and stimuli

Participants completed eleven runs of echo-planar imaging in a single session. All of the functional MRI and behavioural results reported in the present study were acquired in the first run of scanning. The remaining ten runs of scanning, from Run 2 to Run 11, were designed for a separate project, and those data are not reported here. In the present study, we focused on the oddity detection experiment in which participants were required to identify either a semantic anomaly or a geometric anomaly from an array of four items. As illustrated in Figure 1A, in each trial, we presented a quadruplet of visual stimuli; one of the four was an ‘oddball’ that was either semantically or visuospatially inconsistent with the other three items. We used a 2-by-2 factorial design in which we orthogonally manipulated the types of stimuli (words *vs*. polyominoes) and the extent of difficulty (easy *vs*. hard), yielding four task-conditions. The overarching task requirement was identical in every condition – participants were required to find out the oddity that differed from other items. However, words and polyominoes entailed different cognitive operations to achieve the same behavioural goal of identifying an oddball. Detecting a semantic oddity required comprehending the meaning of each constituent word and deriving an abstract conceptual relationship amongst them, whereas detecting a visuospatial oddity required analysing the visual features of each polyomino and mentally rotating them to derive their spatial relationship when necessary. A session began with the acquisition of a structural scan, followed by the oddity detection experiment (Run 1), and then a separate experiment (Run 2 – Run 11). To sustain uninterrupted engagement in a mental state for sufficient time, we used a block design. The experiment comprised 48 task-episodes (each of the four task-conditions contained 12 blocks; each block was 12-sec long, consisting of three trials), 47 inter-block intervals (each 1.5 sec), and a 1-sec blank at the final moment of the scan, yielding a total duration of 647.5 sec. Each trial began with a fixation cross (0.1 sec), followed by a quadruplet of stimuli (3.9 sec); the quadruplet consisted of either words or polyomino patterns, each bounded inside a square. Participants were instructed to respond as quickly as possible within the time-limit. Stimuli display and response collection were controlled using E-Prime (Psychology Software Tools). We fully counterbalanced the order in which the four task-conditions were presented so that each condition was equally likely to appear in every possible timeslot within the sequence (and each condition was equally likely to precede or succeed any other condition), with stimuli randomly drawn from the designated stimuli-sets and shuffled across blocks. When performing the task, participants reacted by pressing one of the four designated buttons on an MR-compatible response-pad using their right hand. The oddball’s location varied randomly trial-by-trial and was equally probable to appear in any of the four locations. All text stimuli were white colour, displayed on a black background; text stimuli were Arial typeface, 24-point in font size. Stimuli were displayed using an MRI-specialised LCD screen (1,920 × 1,080; NordicNeuroLab) and projected onto a coil-mounted device for viewing.

**Figure 1.**
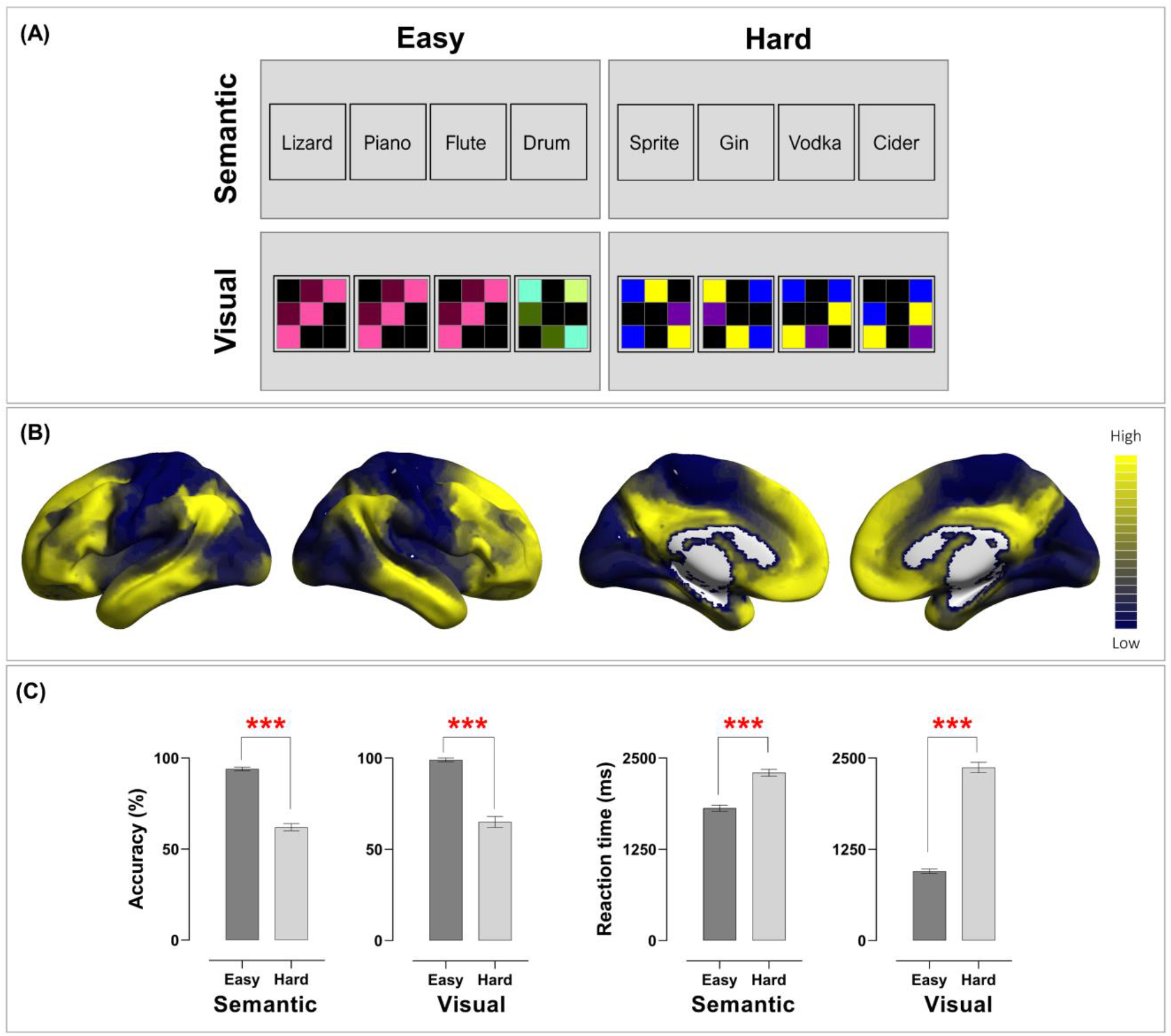
**(A)** Example stimuli of the four conditions. In this 2 × 2 factorial design, the type of cognitive operation (Semantic *vs*. Visuospatial) and the extent of cognitive effort needed to identify an oddball (Easy *vs*. Hard) were independently manipulated. **(B)** The principal gradient that explains the brain’s connectivity pattern, identified by Margulies *et al*. (2016). Brain regions that fall on the topmost tier tend to belong to the default-mode and semantics-related systems, whereas those fall in the lowest tier belong to the sensory-motoric system. **(C)** In both accuracy and reaction times, the difficulty effect was robustly found in both Semantic and Visuospatial conditions

For the semantic anomaly task, we constructed 72 ‘odd-one-out’ quiz questions; 36 questions were designed for the Semantic-Easy condition, while 36 were designed for the Semantic-Hard condition. The complete lists of the 72 questions are reported in *Supplementary Materials*. Each of the questions was presented only once to prevent dwindled neural reaction to repeated exposures. Each question contained a target (semantic oddball) and three foils. Questions of the two conditions differed on the degree of scrutiny required to differentiate semantic concepts. The Semantic-Easy questions were designed based on the following two rules: (***i***) The semantic oddball belongs to a different basic-level linguistic category from the three foils. (***ii***) The three foils are semantically related to each other, while the oddball is unrelated to any of them. With these principles, we constructed a set of questions in which every semantic oddball easily stands out from the quadruplet as it has clearly distinct features from the remaining words. For instance, in the quadruplet of ‘*Lizard, Piano, Flute, Drum*’, all of the three foils belong to the broad category of music instruments and are semantically unrelated to the reptile animals of lizard. The Semantic-Hard questions were designed with the following rules: (***i***) All of the four words in a quadruplet belong to the same basic-level category. (***ii***) The oddball is semantically related to the three foils and only differs from them on an idiosyncratic semantic attribute at a subordinate-concept level. For example, in the quadruplet of ‘*Sprite, Gin, Vodka, Cider*’, all of the four words belong to the broad category of beverages and are ingredients for making cocktails; however, Sprite is the only non-alcoholic drink. To further maximise the disparity between the Easy and Hard condition in the difficulty of semantic retrieval, we deliberately selected words with lower lexical frequency for the Hard condition (hence, slower lexical access and longer retrieval) than the Easy condition (frequncy per million words: Easy - 5,499±1,287, Hard - 526±134, *p* < 0.001; the corpus statistics based on Van Heuven *et al*. 2014). Note, dissociating the effect of linguistic category from lexical frequency was not the aim of our study; therefore, these two factors were compounded so as to maximise the difficulty level between the Easy and Hard condition.

For the visuospatial anomaly task, we created 72 polyomino patterns. These stimuli were used to construct 72 quadruplets (non-repeated); 36 were in the Visuospatial-Easy condition; the remaining were in the Visuospatial-Hard condition. Example stimuli of the two conditions are illustrated in Figure 1A. Each polyomino pattern consists of a black background and 3-by-3 intersecting grids; within a pattern, five cells from the nine positions were tinted with different colours. As shown in Figure 1A, in the Visuospatial-Easy condition, the three foils have exactly identical configuration, orientation, and colours, while the oddball is saliently distinct from all other items (akin to the classic ‘pop-out’ phenomenon in the visual search literature; Treisman and Gelade 1980). By contrast, in the Visuospatial-Hard condition, the three foils have the same configuration and colours, but they are rotated to 0°, 90°, 270° (essentially, they are the same stimulus shown in three different orientations). The oddball in the Visuospatial-Hard condition has a similar visual configuration to its accompanying foils and differs subtly on the spatial arrangement of one or two cells.

### fMRI acquisition

MRI data was collected using a Siemens 3-Tesla PRISMA system. T_1_-weighted anatomical images were acquired using a 3D Magnetization Prepared RApid Gradient Echo (MPRAGE) sequence [repetition time (TR) = 2250 ms; echo time (TE) = 3.02 ms; inversion time = 900 ms; 230 Hz per pixel; flip angle = 9°; field of view (FOV) 256 × 256 × 192 mm; GRAPPA acceleration Factor 2]. Functional task-evoked data were obtained with a multi-echo multi-band (MEMB) blood oxygenation level dependent (BOLD)-weighted echo-planar imaging (EPI) pulse sequence. This MEMB sequence has the strengths that it acquired four functional volumes for each TR (multi-echo, enabling capturing signals that peaked at early and late echo-times that are often overlooked by conventional protocols) and simultaneously recorded two slices during the acquisition of each volume (multi-band, speeding up the acquisition rate). The parameters included: TR = 1,792 ms; TE_1_ = 13 ms, TE_2_ = 23.89 ms, TE_3_ = 34.78 ms, TE_4_ = 45.67 ms; flip angle = 75°; FOV = 192 mm × 192 mm, MB-Factor = 2, in-plane acceleration = 3. Each EPI volume consisted of 46 axial slices in descending order (3 × 3mm; starting concomitantly from the top and middle slice) covering the whole brain (FOV = 240 × 240 × 138 mm). For the present study, a series of 362 functional volumes were acquired for the oddity detection task.

### Pre-processing

All raw DICOM data were converted to NifTi format using dcm2niix. The T_1_ structural images were processed using the standard processing pipeline of the FSL package’s ‘fsl_anat’ function (Ver5.0.11; https://fsl.fmrib.ox.ac.uk/). This pipeline involves these sequential six steps: (***i***) Reorient images to standard MNI space (‘fslreorient2std’), (***ii***) automatically crop image to remove the neck (‘robustfov’), (***iii***) bias-field correction to fix field inhomogeneity (‘fast’), (***iv***) registration into the MNI space (‘flirt’ and ‘fnirt’), (***v***) brain extraction (using ‘fnirt’ nonlinear method) and (***vi***) tissue-type segmentation to separate white-/grey-matter and other structures (‘fast’). Each T_1_-image was individually inspected for accuracy after being normalised into the MNI space. The functional EPI data were pre-processed using a combination of tools in FSL, AFNI (Ver18.3.03; https://afni.nimh.nih.gov/), and a specialised Python package to perform TE-dependent analysis (Kundu *et al*. 2012; Kundu *et al*. 2013; Kundu *et al*. 2017). Despiked (‘3dDespike’), slice-time corrected (‘3dTshift’, matched to the middle slice), and realigned (‘3dvolreg’) images were submitted to the “tedana” toolbox, which took the time-series data from all of the four TEs and decomposed the resulting data into BOLD-signals and noises (non-BOLD components). Specifically, decomposition was based on whether a signal series depended on the four echo-times – with the strength of multiple echo-times, the algorithm was able to tell apart noises that fluctuated randomly or independently of the timings of four TEs (e.g., scanner’s drift, cardiac/respiratory noises, head motion) from signals that systematically varied with the timings of TEs (e.g., the functional data of BOLD). Data of the four echo-times were then optimally integrated, weighted based on the intensity of T_2_* signal in each voxel and separated from the TE-independent/ non-BLOD noises (Kundu *et al*. 2017). Finally, the optimally-combined images were co-registered into each individual’s T_1_ structural scan (using FSL’s ‘flirt’), normalised to the standard MNI space (using FSL’s ‘fnirt’ warps and ‘flirt’ transform), and smoothed with a 6 mm FHWM Gaussian kernel.

### General linear model and psychophysiological-interaction connectivity

The SPM12 package (https://www.fil.ion.ucl.ac.uk/spm/software/spm12/) was used to construct a general linear model (GLM) for subsequent analyses. For each participant, the onset times and duration of every task-episode were used to create an experimental-design matrix. Each individual’s design matrix was convolved with a canonical haemodynamic response function. We included each participant’s reaction times (RTs) as a parametric modulator that were attached to each task regressor, allowing us to take into account neural activities driven by task difficulty or cognitive effort when assessing the effects of experimental manipulation. Contrast images from the fixed-effect model at individual-level (1^st^-level) were submitted to the random-effect model at the group-level (2^nd^-level). We statistically thresholded the whole-brain interrogation GLM results at *p* < 0.05 (FWE-corrected for multiple comparisons) for clusters and *p* < 0.001 for voxel intensity.

To investigate how context-dependent connectivity to the inferior frontal gyrus (IFG) altered between linguistic and non-linguistic situations, we used SPM12 to perform a psychophysiological-interaction (PPI) analysis. The IFG, also known as the Broca’s area, has been implicated in a broad range of language-related processes (for review, Fedorenko and Blank 2020), particularly when participants allocated greater amount of cognitive resources to solve a semantic problem (e.g., Chiou *et al*. 2018). We used the left IFG as a region of interest (ROI) and individually defined its locus for each person, guided using the group-level peak activation coordinate (*x* = -44, *y* = 24, *z* = -2) from the contrast of ‘Semantic-Hard > Semantic-Easy’ (which identified brain regions associated with higher difficulty of semantic processing). For each individual, we pinpointed the (Semantic Hard > Easy) local maxima of IFG activation nearest to the group-level peak coordinate (searched within the scope of 8mm radius) and set it as the ‘seed’ of PPI connectivity. At each individual’s IFG peak, we created a spherical ROI (radius = 6mm) and extracted the first eigenvariate in the sphere using SPM12’s built-in algorithm. The eigenvariate was a proxy of the seed’s underlying neural/physiological activities. It was then convolved with the psychological factor (the contrast of cognitive states: Semantic *vs*. Visuospatial). This generated the interaction term – the psychophysiological/PPI factor that denoted changes in connectivity with the IFG seed as a function of switching between task-conditions. These three factors – the psychological, physiological, and PPI – were used to construct a GLM for whole-brain search. We focused on the PPI regressor to identify brain regions whose neural connectivity with the left IFG was modulated as a function of Semantic *vs*. Visuospatial context. Statistical thresholds were the same as those for the GLM analysis – *p* < 0.001 (voxelwise) and *p* < 0.05 (FWE-corrected for clusters).

### Cortical gradient analysis

To understand how the neural activity triggered by different task-conditions were couched within the macroscale architecture of whole-brain connectivity, we projected various fMRI results onto the principal (first) gradient of Margulies *et al*.’s (2016) hierarchical framework of brain organisation. The methodological details of deriving the cortical gradients were reported in the original study. Here we summarised their main analytical steps: Using the resting-state fMRI data of 820 participants from the Human Connectome Project combined with different nonlinear dimension-reduction methods (e.g., Laplacian eigenmaps and diffusion map embedding), Margulies *et al*. analytically reduced the complexity of connectivity into gradients that succinctly delineated the majority of variance regarding how regions are functionally linked together. The principal gradient, which explained the greatest variance, was anchored, at one end of the spectrum, by primary sensory-motoric regions that directly receive input from the external world or generate a response to interact with the environment; at the other end of the spectrum, the gradient was anchored by default-mode regions that are involved in abstract cognition (see Figure 1B). The original gradient structure assigned each voxel of the brain a value between 0 and 100, relating to where it fell on the gradient (0 = sensory-motor end; 100 = default-mode end). For our analysis, this gradient was divided into 20 bins, with voxels within each five-percentile bin lumped together (for example, all voxels with a value between 0 and 5 were grouped in Bin-1, and all voxels ranged 6 – 10 were grouped within Bin-2, etc.; each bin contained nearly identical number of voxels – range: 6,431 – 6,441; mean±SD: 6,436±2) (for precedents of this approach, see Murphy *et al*. 2018; Murphy *et al*. 2019; Lanzoni *et al*. 2020; Wang *et al*. 2020). Next, we used each of the 20 five-percentile bins as a region of interest (ROI) and extracted activation amplitudes (from GLM) and connectivity strengths (from PPI), allowing us to investigate how the neural correlates were distributed across the gradient. The original gradient data of Margulies *et al*. (2016) are publicly available online (https://www.neuroconnlab.org/data/index.html).

## Results

### Behavioural performance

The overarching behavioural goal was *identical* in every task-condition (identifying an oddball from a quadruplet) but the stimuli and required operations differed between tasks. Figure 1C illustrates the group-level results of accuracy rates and reaction times. To ascertain whether our tasks effectually modulated the difficulty of semantic and visuospatial processing, we performed *a priori* tests to compare performance under the Easy and Hard condition for the Semantic task and Visuospatial task, respectively. The effects were reliably detected in both tasks – in the Semantic task, identifying an semantic oddball was more accurate (*t*_(24)_ = 13.91, *p* < 0.001) and quicker (*t*_(24)_ = -13.70, *p* < 0.001) in the Semantic-Easy than Semantic-Hard condition. Similarly, in the Visuospatial task, identifying a perceptual oddball was also more accurate (*t*_(24)_ = 14.08, *p* < 0.001) and quicker (*t*_(24)_ = -21.34, *p* < 0.001) in the Visuospatial-Easy than Visuospatial-Hard condition. These results confirmed the efficacy of our difficulty manipulation for both tasks and warranted our subsequent search for effects at the neural level. Next, we performed a 2-by-2 repeated-measure ANOVA, with Task and Difficulty being within-participant variables. In reaction times, there was a significant interaction between Task and Difficulty (*F*_(1, 24)_ = 217.38, *p* < 0.001, *η*^2^_p_ = 0.90). Post-hoc tests were performed to dissect the source of this interaction – while the Semantic-Hard and Visuospatial-Hard conditions yielded comparable reaction times (*p* = 0.28, *n*.*s*.), reaction times were faster in the Visuospatial-Easy than in Semantic-Easy condition (*p* < 0.001). The same 2×2 interaction was not significant in the accuracy data (*F* < 1). Taken together, the behavioural results suggest that our experimental design effectively induced robust difficulty effects in both semantic and visuospatial domains. Following the precedents of Mineroff *et al*. (2018), we used (Visuospatial-Hard > Visuospatial-Easy) to define MD regions and (Visuospatial-Easy > Visuospatial-Hard) to define default-mode regions. Semantic control regions were defined by (Semantic-Hard > Semantic-Easy), and the language network by (Semantic-Hard > Visuospatial-Hard, owing to their difficulty level being equated).

### Topographical alignment of the multi-demand, semantic-control, and default-mode systems

We began by employing traditional group-level analysis to examine the neural activity triggered by semantic control and compared the topographical location of this network with MD regions and default-mode regions. As illustrated in Figure 2(A), whole-brain interrogation based on ‘Vis.-Hard > Vis.-Easy’ revealed a set of frontoparietal areas that resembled a typical topography of the MD system (the blue inset box: the posterior dorsolateral frontal cortex, intraparietal sulcus, insula, middle/anterior cingulate cortex), while the reverse contrast of ‘Vis.-Easy > Vis.-Hard’ detected a group of widely dispersed regions that constituted the default-mode system (the green inset box: the rostro-medial prefrontal cortex, angular gyrus, posterior cingulate cortex, and middle temporal gyrus). Critically, semantic control (Sem.-Hard > Sem.-Easy; the red inset box) triggered a set of areas that are, topographically speaking, sandwiched between the MD network and default-mode network, giving rise to an orderly pattern seen in multiple sections of the cortex. To highlight the intermediary status of semantic control areas, we present the medial prefrontal cortex and the left lateral-parietal cortex as examples in Figure 2(A). As clearly illustrated in 2(A), there was an orderly transition in the medial prefrontal cortex, progressing from an MD area (blue: anterior cingulate cortex), through semantic control in the middle (red: dorsomedial prefrontal cortex), to a default-mode cluster in the most anterior subpart (green: rostromedial/ventromedial prefrontal cortex). Also in Figure 2(A), a similar ordering was observed in the lateral parietal cortex, from MD regions in the superior parietal lobule and intraparietal sulcus (blue), via a semantic control patch in the middle (red), to default-mode regions in the angular gyrus (green). This orderly arrangement was also observed in the left IFG (see *Supplementary Results 1*), shifting from the posterior IFG that preferred visuospatial control (*pars opercularis*: MD cluster), through the middle-to-anterior zone preferring semantic control (*pars triangularis* and *pars orbitalis*), to the most anterior part of IFG that preferred default-mode processes (*pars orbitalis* and frontal pole). Also shown in *Supplementary Results 1*, there was a graded transition in the occipital/temporal lobes, from expansive MD clusters in the posterior-inferior occipitotemporal regions, through semantic control in intermediate zones (the pMTG), to default-mode clusters in the anterior-superior temporal regions. In summary, in all four of these left-hemisphere cortical segments that we inspected, semantic control was situated in intermediate positions, abutted by MD clusters and default-mode clusters from two sides.

**Figure 2.**
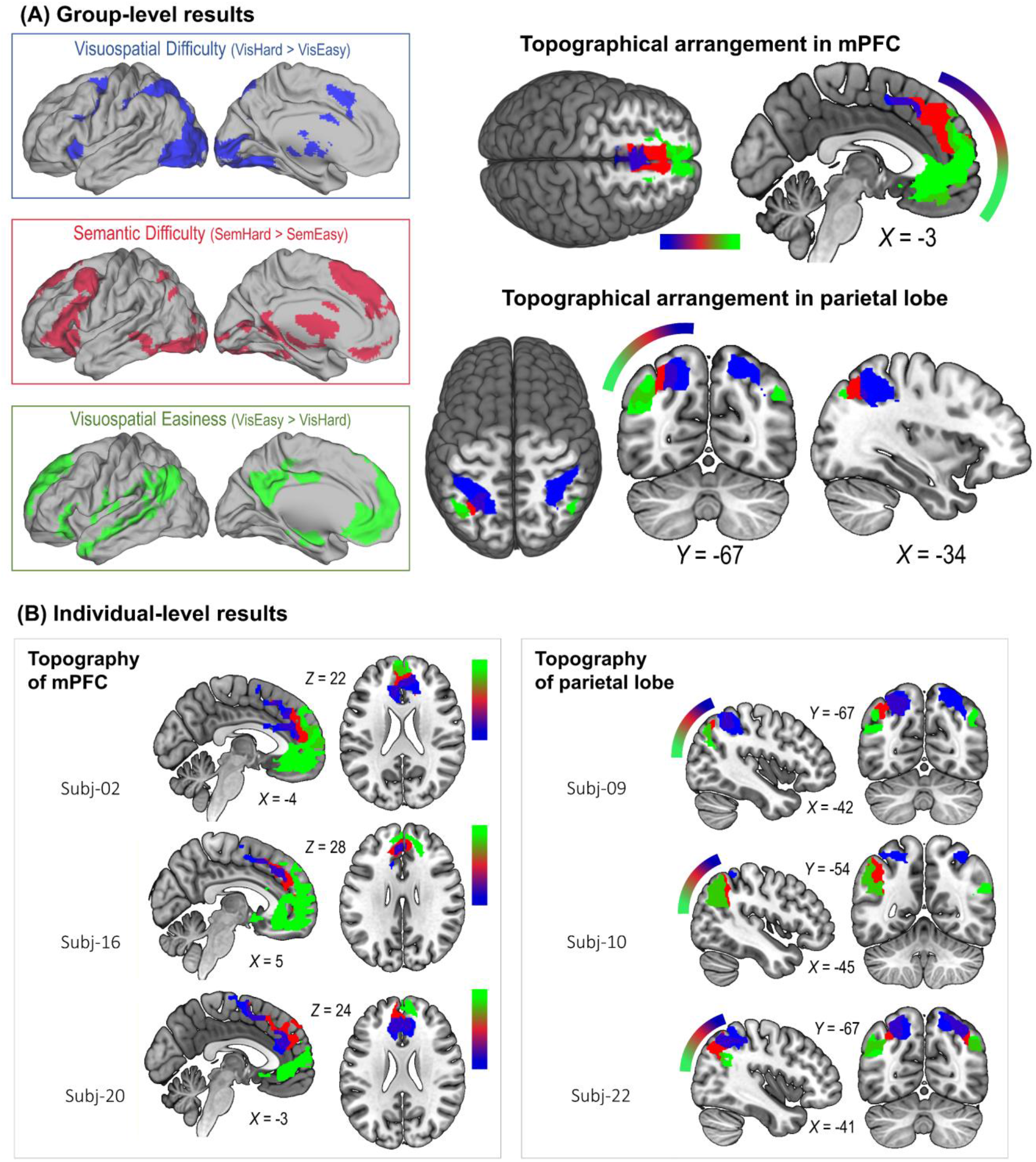
**(A)** The inset boxes show the whole-brain search for the effects of visuospatial difficulty (blue), semantic difficulty (red), and visuospatial easiness (green). The Suprathreshold clusters of the three effects were found to form orderly arrangements in various parts of the brain – shown here are examples in the medial prefrontal cortex and lateral parietal cortex lobe. Statistical threshold of group-level analysis: *p* < 0.001 at the voxel-wise level, *p* < 0.05 (FWE-corrected) at the cluster level. **(B)** Similar orderly arrangements (wherein semantic control was flanked by visuospatial difficulty and visuospatial easiness) were also found in the individual participant results.

A problem with group-level analysis is its lack of ability to accommodate individual variation and loss of precision (for discussion, see Fedorenko 2021). In our case, while group-level results revealed an orderly pattern, the loci of semantic control activation might have varied across individuals, with only voxels in intermediary loci consistent enough in group-level results. To ascertain whether the orderly pattern could also be identified at the individual level, we scrutinised the topography of every participant and found a highly consistent pattern across the group – although the exact loci of MD, semantic control, and default-mode clusters somewhat differed among volunteers, the topography in an individual brain reliably emerged as a layout wherein semantic control was juxtaposed between MD and default-mode. In Figure 2(B), we illustrated the results of six example participants – while the precise positions of clusters and their extent of overlap varied between participants, the clusters of semantic control reliably occupied the intermediate expanse of cortex, flanked by MD voxels and default-mode voxels. This topography was consistently found across the group, and the unthresholded group-level statistical maps (semantic control, MD, and default-mode results) have been uploaded to NeuroVault for interested readers to examine their cortical distribution. Taken together, these results echo previous evidence about the topography of the MD system that this domain-general system adjoins nearby regions that have domain-specific tuning (e.g., regions referentially react to auditory or visual tasks, see Assem *et al*. 2021; in the present study, MD areas abuts semantic-control areas). Furthermore, our data raise an interesting possibility that the functional specialty of a brain region in the association cortex might be influenced by the areas that encircle it.

### Situating semantic control *vs*. visuospatial control along the principal cortical gradient

We projected the unthresholded whole-brain reaction to semantic difficulty (Sem.-Hard > Sem.-Easy) onto the principal gradient, evaluated its distribution across 20 percentile bins from sensorimotor to heteromodal cortex, and compared its trajectory with visuospatial difficulty (Vis.-Hard > Vis.-Easy). Figure 3 shows that the maximal reaction of visuospatial control was situated in lower/middle portions of the gradient (Tier 3-12, shaded with blue), whereas the response of semantic control gradually ramped up along the gradient and peaked at the topmost bin (Tier 15-20, shaded with orange). ANOVA identified a significant interaction of gradient locations (20 bins) by the two types of control (semantic *vs*. visuospatial): *F*_(19, 456)_ = 48.43, *p* < 0.001. Thus, post-hoc tests were performed to identify the source of interaction: From Tier 3 to 12 (blue range, near the primary sensorimotor cortex), cortical areas showed greater response to visuospatial difficulty than semantic difficulty (all *p* < 0.05). From Tier 15 to 20 (orange range, encompassing the most heteromodal zones), cortical areas showed greater activation to semantic difficulty than visuospatial difficulty (*p* < 0.05 in all bins of this range). The gradient results informed where the two types of control are situated the process of integration: Semantic control was operated on abstract representations in heteromodal (highly integrated) zones, whereas visuospatial control was operated on modality-specific, less integrated representations.

**Figure 3.**
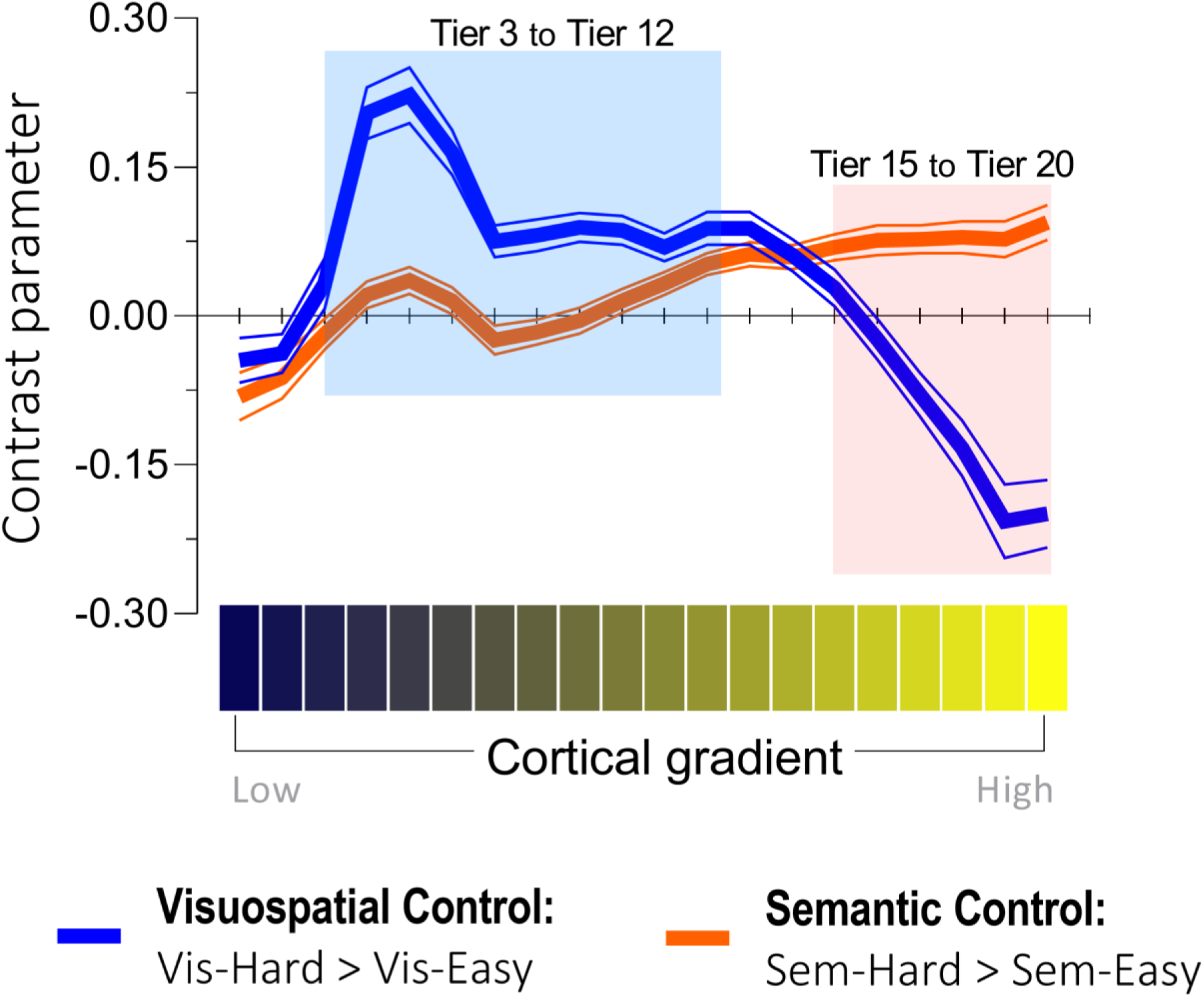
The effects of visuospatial control (blue curve) and semantic control (orange curve) were projected onto the 20 percentile tiers of the principal gradient. A significant interaction was detected between the 20 gradient tiers and the two types of control. Based on the interaction, post-hoc tests were performed to compare the difference between the two control processes in each bin. Cortex in Tier 3 to 12 (shaded in blue) was more active for visuospatial control than semantic control, whereas cortex in Tier 15 to 20 (shaded in orange) showed the opposite pattern.

### The interaction effect in semantic control regions: *semantic difficulty* and *visuospatial easiness*

We performed a whole-brain interrogation to identify regions that exhibited a significant interaction: The interaction was defined as *semantic difficulty* (Sem.-Hard > Sem.-Easy) and *visuospatial easiness* (Vis.-Easy > Vis.-Hard). Note, this analysis computed the *interaction term* (opposite pattern in the semantic and visuospatial tasks), rather than simply aconjunction of the two contrasts. This interaction opposite preferences for difficulty in different domains – enabled a stringent test to identify regions that showed a domain-specific proclivity for semantic difficulty, rather than a generic preference for all kinds of difficult operations. Specifically, we examined (***i***) which regions exhibited this interaction; (***ii***) in the post-hoc analysis, whether the regions that showed the interaction were significantly more active for difficult semantic decisions than for easy visuospatial decisions.

As illustrated in Figure 4(A), the interaction analysis revealed (***i***) a group of areas well-established in the semantic control literature (the left IFG, left pMTG, and dmPFC; see Jackson 2020 for a review of regions involved in semantic control), and (***ii***) a group of areas within the default-mode network (e.g., the posterior cingulate cortex, ventromedial prefrontal cortex, and inferior parietal lobule, etc.)^1^. Next, we investigated the distribution of this interaction along the principal gradient. As shown in Figure 4(B), areas situated towards the heteromodal end of the gradient were increasingly more active for ‘semantic difficulty and visuospatial easiness’. By contrast, areas towards the sensorimotor end of gradient showed the *opposite* interaction: ‘visuospatial difficulty and semantic easiness’. While the whole-brain analysis above revealed regions showing a significant interaction between difficulty and domain, it did not specify the details of this interaction – namely, whether this region reacted equally to semantic difficulty and visuospatial easiness, or whether it had a larger effect of semantic difficulty than visuospatial easiness (or the opposite pattern). To clarify this, we examined the activation pattern at the cluster peaks (i.e., this procedure was akin to conducting *post-hoc* tests following a significant interaction). For the nine suprathreshold clusters that fulfilled this interaction, we created a spherical ROI (6-mm radius), centred at the peak of each cluster, and extracted the parameters for the contrasts of ‘Sem.-Hard > Sem.-Easy’ and ‘Vis.-Easy > Vis.-Hard’. As shown in Figure 5, many areas of the default-mode system preferred visuospatial easiness to semantic difficulty. By contrast, the IFG and dmPFC, two areas known for their involvement in semantic control, showed significantly larger responses for semantic difficulty than visuospatial easiness^2^. Taken together, these analytical procedure (interaction, followed by post-hoc tests) reaffirmed the unique profile of semantic control regions – they were *not* triggered by visuospatial difficulty (instead, they preferred visuospatial easiness) and their difficulty-driven responses were tuned to semantics.

**Figure 4.**
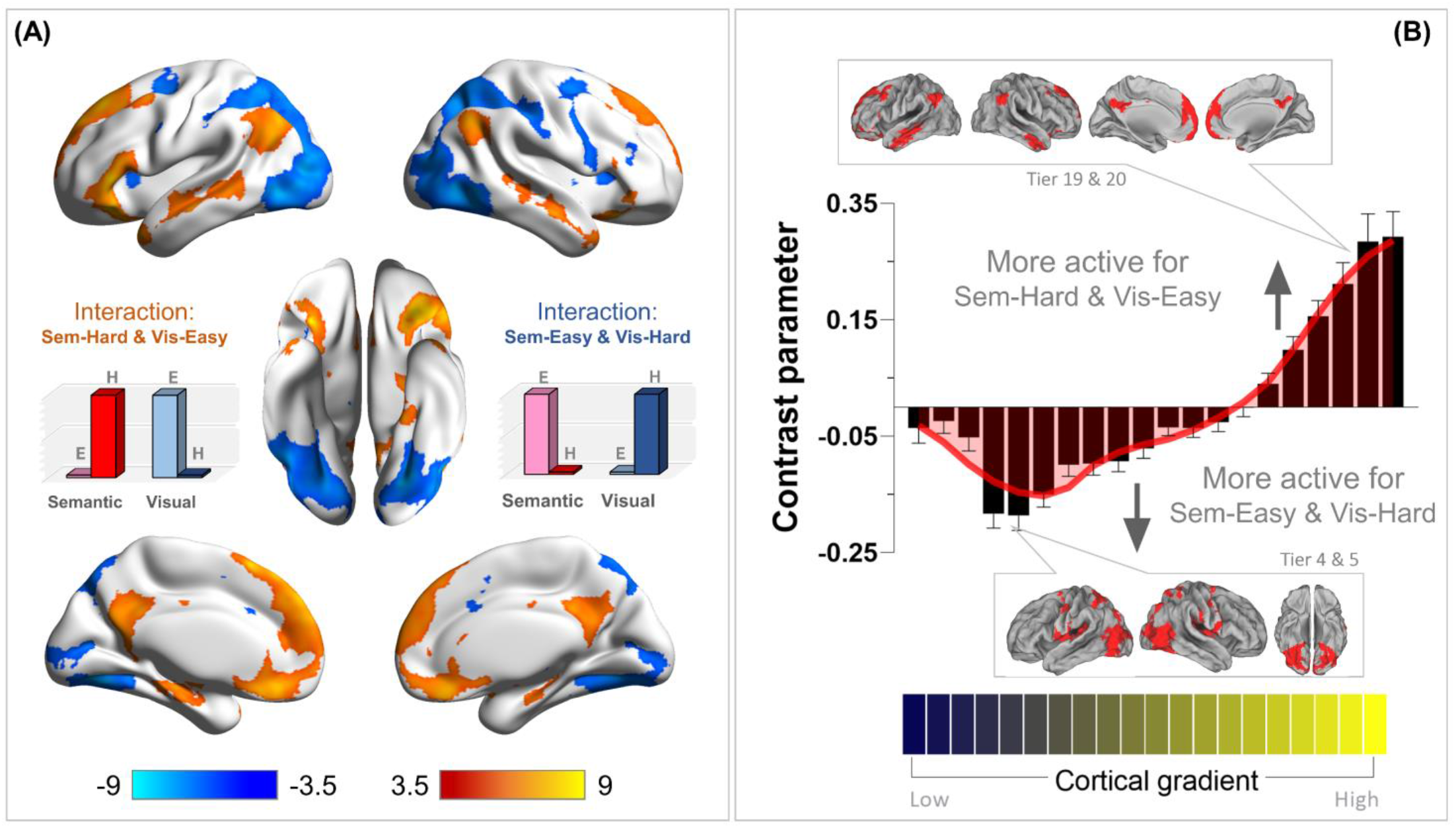
**(A)** Whole-brain search for the interaction of ‘semantic difficulty and visuospatial easiness’ (in ward colour) and the interaction of ‘visuospatial difficulty and semantic easiness’ (in cold colour). Statistical threshold of group analysis: *p* < 0.001 at the voxel-wise level, *p* < 0.05 (FWE-corrected) at the cluster level. **(B)** Projecting the interaction onto the 20 bins of the cortical principal gradient. The gradient tiers in which the maximal interaction effects occurred were highlighted in the insets.

**Figure 5.**
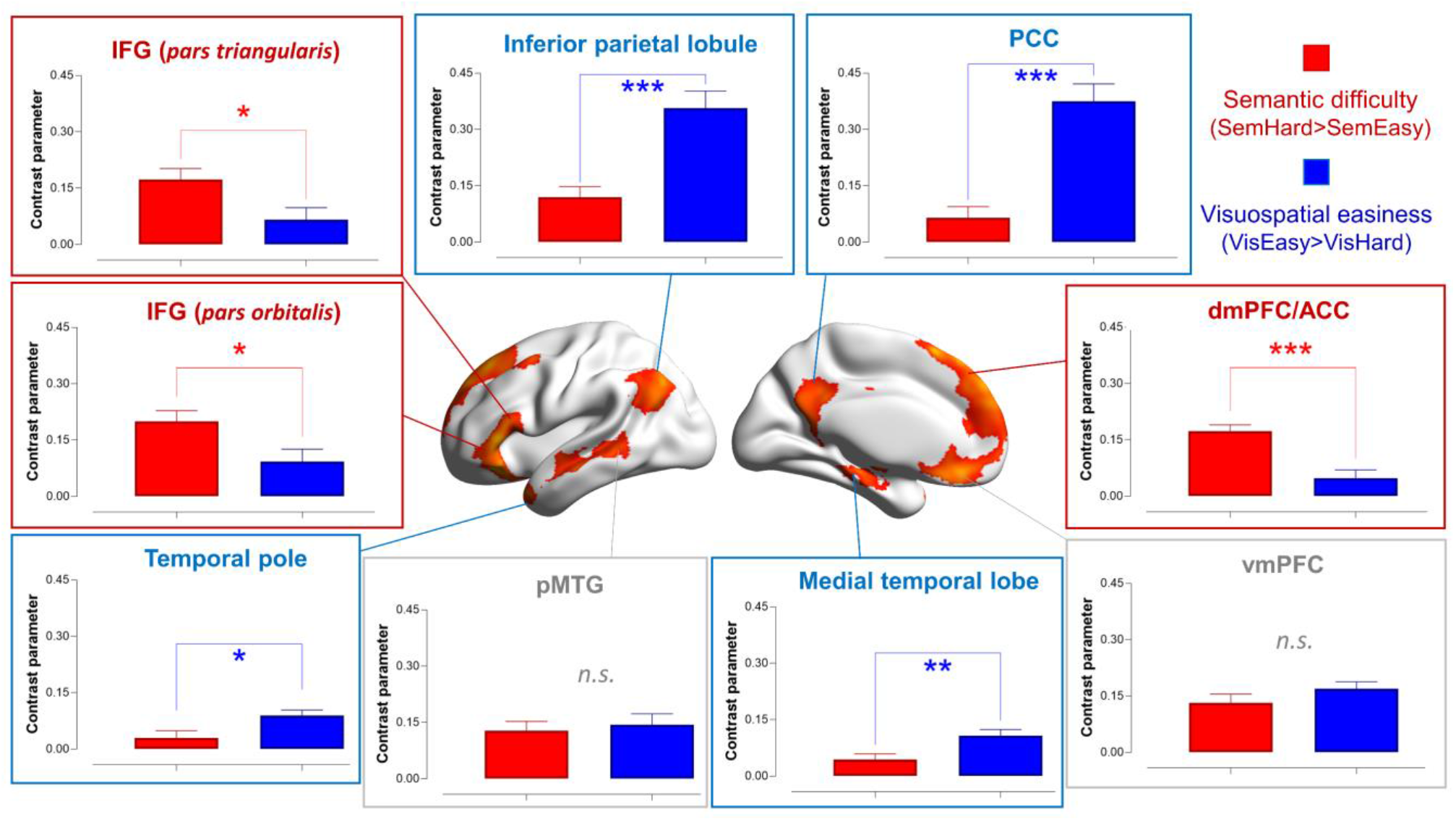
Based on the significant interaction, post-hoc comparisons were conducted at the peaks of the nine clusters showing a significant interaction of ‘semantic difficulty and visuospatial easiness’.

### PPI analysis on context-reliant connectivity to the IFG and its distribution along the gradient

The left IFG, a region known for its contribution to semantic cognition, has an intriguing profile in its reaction to different task situations and affiliation with different networks. Above we show that the left IFG exhibited an interaction (semantic difficulty and visuospatial easiness; Figure 5). Thus, unlike the MD system that invariably prefers harder to easier operations under diverse situations, the IFG is selectively tuned to cognitive effort for semantic tasks. This concurs with previous literature that IFG activation elevated for difficult semantic tasks (Chiou *et al*. 2018) and reduced for difficult perceptual task (Chiou *et al*. 2020). Connectivity-wise, the IFG in the intrinsic state is linked to the default-mode network (Yeo *et al*. 2011); however, it is also connected with semantics-related regions of the language network (Jackson *et al*. 2015) and some sections of the multiple-demand network (Humphreys and Lambon Ralph 2017). It is one of few brain regions that form functional and structural connectivity with both the multiple-demand network (e.g., with the inferior frontal sulcus) and with the semantic network (Davey *et al*. 2016). These results suggest that the left IFG might serve as a ‘switchboard operator’ mediating the conversation between different specialised systems, which gives rise to its mixed functional profile. To further explore this, we performed a PPI analysis to examine how neural connectivity with the left IFG altered as a result of our experimental contexts.

As illustrated in Figure 6(A), we placed the seed for this connectivity analysis at each individual’s IFG peak activity for the ‘semantic difficulty’ contrast; whole-brain analysis identified voxels whose connectivity with the seed varied between the Semantic and Visuospatial contexts. We found that during the Semantic condition, the left IFG was more tightly coupled with the bilateral MD network and visual cortex; by contrast, during the Visuospatial condition, it was more connected with the default-mode system. This pattern of PPI-connectivity is the exact antithesis of what the contrast of task-related activities revealed (see the inset box of 6A): the MD system and visual cortex were more active for the Visuospatial task, while the default-mode system and language-related areas were more active for the Semantic task. These results suggest that when the task requires an interface between perception (i.e., text stimuli) and semantic operation (i.e., analysing conceptual links between words to discover semantic oddity), the left IFG played an intermediary role to facilitate the communication between dorsal frontoparietal regions (i.e., the MD system) and ventral frontotemporal regions (i.e., the language system; Davey *et al*. 2016). However, when the cross-talk between systems was not necessary (such as during resting-state or during the Visuospatial task in which the stimuli conveyed no semantic meaning), the left IFG was aligned with regions preferentially tuned to memory processes (i.e., default-mode system or language-related areas), consistent with the resting-state literature (e.g., Yeo *et al*. 2011).

**Figure 6.**
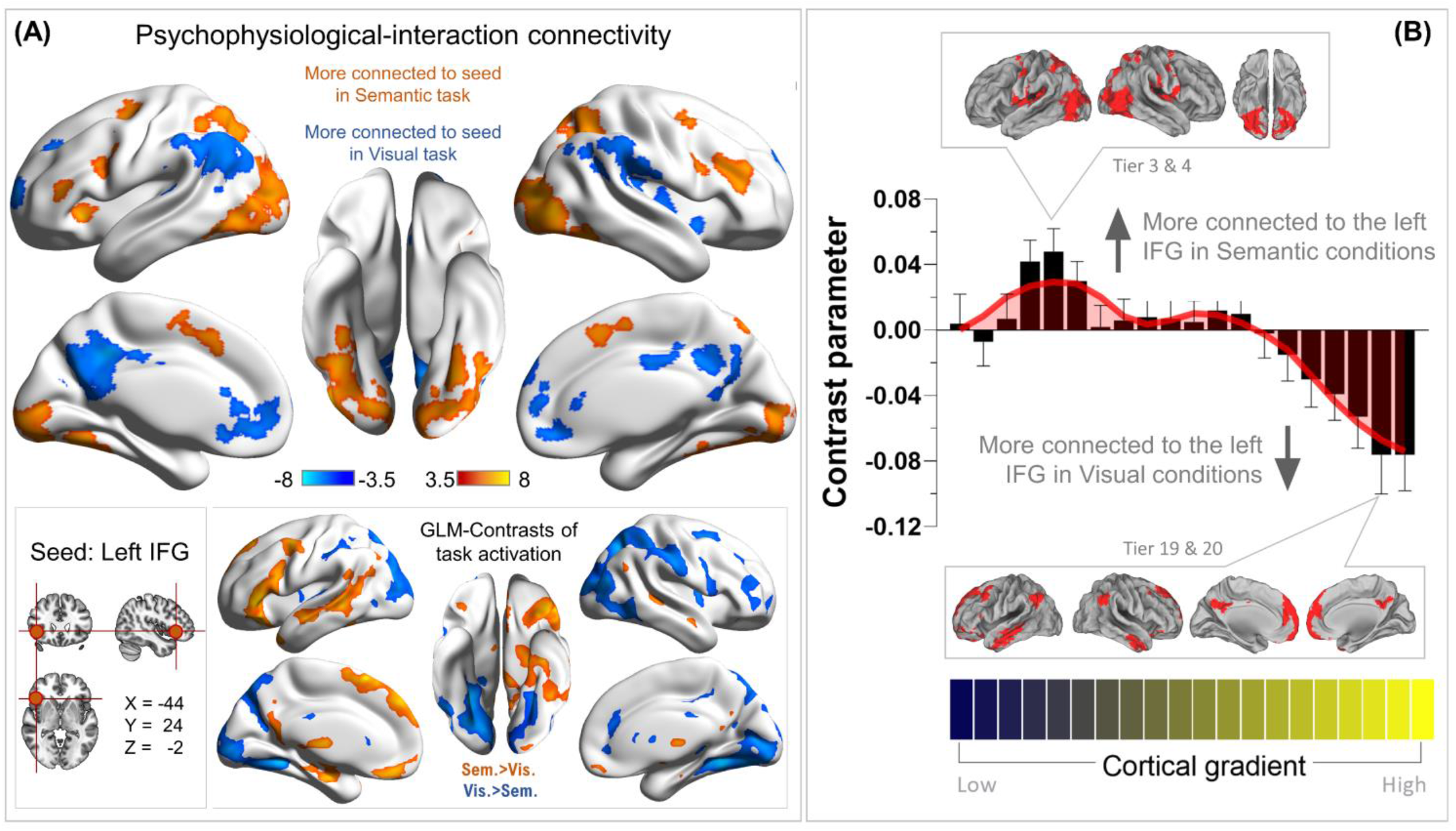
**(A)** The PPI results of contextually dependent connectivity to the seed region of left IFG, comparing how the connectivity pattern altered between the Semantic and Visuospatial conditions. Statistical threshold of group analysis: *p* < 0.001 at the voxel-wise level, *p* < 0.05 (FWE-corrected) at the cluster level. **(B)** The PPI results were projected onto the principal gradient of the human brain. The gradient tiers in which the maximal PPI effects occurred were highlighted in the insets.

Projecting the whole-brain PPI parameters onto the brain’s principal organisational gradient offered further evidence for an intermediary role for left IFG. As Figure 6(B) shows, the sensorimotor end of the continuum (specifically, Tiers 4-6, including many areas of the ‘dorsal-attention’ network) was more connected with the IFG-seed during the Semantic task. By contrast, the ‘heteromodal’ end of the gradient (particularly the three topmost bins that contain mostly areas of the default-mode network) were more connected with the IFG during the Visuospatial task. This pattern of distribution is consistent with the interpretations that (***i***) there was intensified dialogue between the left IFG and MD network (and many regions of the visual cortex) when the context necessitated deriving semantics from perceptual input; (***ii***) while the MD system supports a wide range of cognitively laborious tasks, semantic control areas were recruited to interface executive control with semantic representations.

### Parametric modulation effects that reveal domain-general and domain-specific mechanisms

In addition to the four task conditions used as regressors in the GLM (which allowed us to examine using the categorical/discrete contrasts between conditions), reaction times (RTs) were also included as parametric modulators, allowing us to evaluate the impact of RT fluctuation on neural activation. Here, the regression model estimated the net effect of RT *after* controlling for the effect of condition. This analysis revealed four different aspects of control machinery (Figure 7A): (***1***) *Domain-general amplification*: activation of the MD network intensified with *longer* RTs irrespective of tasks; the domain-general amplification was seen in the middle frontal gyrus, insular cortex, frontal eye field, bilateral posterior/superior parietal lobules, and the visual cortex. (***2***) *Domain-general suppression*: two midline structures of the default-mode network – the medial prefrontal and posterior cingulate cortices – became more active when RTs were *shorter* irrespective of tasks, suggesting that these midline default-mode ‘cores’ generally prefer automatic (easier) to effortful (harder) processes. (***3***) *Specificity to semantic control*: the left IFG and its neighbouring middle frontal gyrus were significantly more positively modulated by semantic RTs than visuospatial RTs, suggesting these regions’ preference for devoting more cognitive effort to tackle semantic than visuospatial difficulty. *Specificity to visuospatial control*: the dorsal-attention network (the bilateral superior/posterior parietal lobules) and visual cortex were more modulated by visuospatial RTs than semantic RTs, suggesting these areas’ sensitivity to greater difficulty in visuospatial domain than semantic domain. Taken together, the parametric analysis highlighted the two facets of the brain’s control mechanisms domain-generality and domain-specificity, complementing our argument that semantic control regions and MD regions were sensitive to distinct types of behavioural signatures.

**Figure 7.**
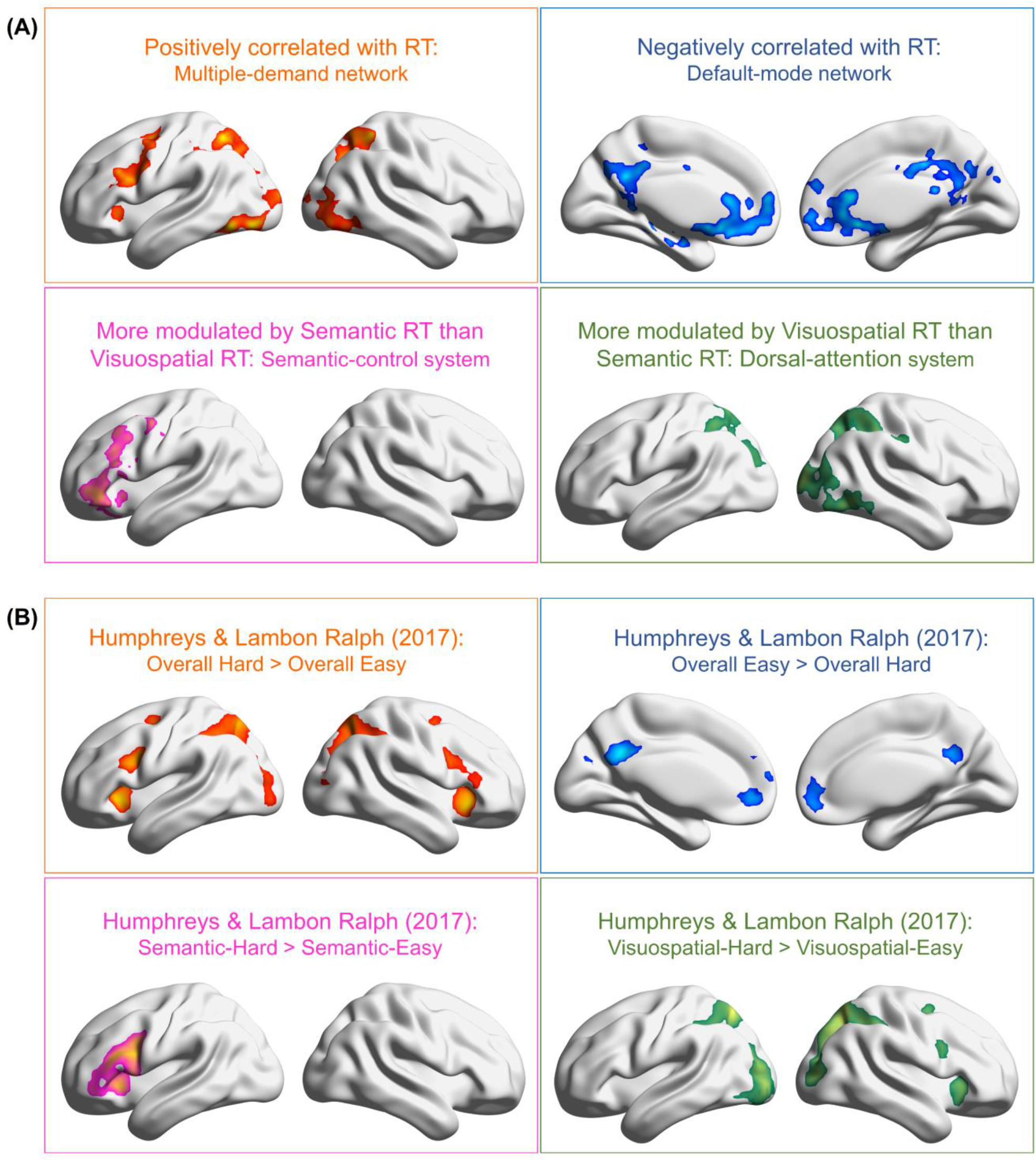
**(A)** Parametric modulation analysis of reaction times revealed four aspects of the brain’s control-related machineries: *domain-general positive correlation with RTs* (multi-demand network), *domain-general negative correlation with RTs* (default-mode network), *semantic control* (the left IFG and adjoining regions), and *visuospatial control* (dorsal-attention system). **(B)** A strikingly consistent pattern of brain activation was replicated using the independent dataset of Humphreys and Lambon Ralph (2017).

We revisited the dataset of Humphreys and Lambon Ralph (2017), qualitatively compared the topography of this previous study with the current findings, and found a strikingly consistent pattern of domain-generality and domain-specificity. In this previous study, participants were required to either judge semantic relatedness or compare visual shapes, and difficulty level (easy *vs*. hard) was manipulated for each task. As shown in Figure 7(B), in the domain-general contrasts of easy *vs*. hard, areas of the MD network were significantly more active for the hard conditions across tasks, whereas areas of the default-mode network were significantly more active for the easy conditions^3^ across tasks. In the domain-specific contrasts, the left IFG was more active for semantic demands, whereas the dorsal-attention system was more active for visuospatial demands. A highly robust pattern was found that replicated across analyses and two independent datasets – the domain-general machinery (invariant to different stimuli/tasks) was supplemented by domain-specific machinery to tackle the challenges uniquely required by the situation.

### Supplementary analyses for the MD system and the language system

In *Supplementary Materials*, we performed additional analyses on the MD and language systems. Specifically, in *Supplemental Results 2*, we parcellated the MD system into subregions based on both hemisphere and atlas demarcation (Tzourio-Mazoyer *et al*. 2002; Fedorenko *et al*. 2013) and tested how semantic- and visuospatial-specific difficulty effects manifested in each subregion. This analysis revealed the anticipated effect that semantic control has larger effects in left-lateralised ROIs whereas visuospatial control was bilateral. Also in *Supplemental Results 2*, we compared the activation pattern for semantic difficulty with meta-analyses data (using NeuroSynth), situating semantic control effects in our task within the maps for ‘semantics’ and ‘executive-control’. This analysis allows a comparison between the present results and relevant literatures. Next, in *Supplemental Results 3*, we report the contrasts of Semantic-Hard *vs*. Visuospatial-Hard because these two conditions had statistically matched behavioural performance levels. The contrast of ‘Sem.-Hard > Vis.-Hard’ revealed a pattern of activation highly similar to the topology of the language-specific system (e.g., Scott *et al*. 2017), whereas the reverse contrast (Vis.-Hard > Sem.-Hard) showed a pattern similar to the MD system (Fedorenko *et al*. 2014).

## Discussion

In the present study, we investigated the cortical topography of semantic control areas in relation to the MD and default-mode systems across multiple analyses: *First*, in the group-level statistical maps, we found that semantic control areas were topographically ‘sandwiched’ between the MD system and the default-mode system. *Second*, we verified the consistency of this pattern by scrutinising the individual-level topographies and found that this orderly pattern could be reliably replicated in individual data (despite the exact loci and cluster-defining thresholds differing between individuals). Topography-wise, semantic control clusters were flanked by the MD and default-mode systems; while some overlap with nearby MD or default-mode clusters could be seen at the fringe, much of the middle territory were recruited by semantic control. Function-wise, semantic control areas showed heightened responses for semantic difficulty and visuospatial easiness (but significantly more active for the former). Such a specific tuning to difficult semantic operation implies a fusion between the tuning of MD regions (tuned to greater difficulty regardless of tasks; Duncan 2010) and default-mode regions (tuned to memory regardless of difficulty level; Smallwood, Bernhardt, *et al*. 2021). *Third*, we projected the activation of semantic control onto the principal gradient (Margulies *et al*. 2016) and found it was couched in different sectors of the gradient from visuospatial control. *Fourth*, the PPI-connectivity evidence showed that the left IFG is a key site that mediated the crosstalk between areas involved in visual perception (dorsal-attention network and visual cortex) and abstract cognition (the semantic and default-mode networks). *Finally*, parametric modulation analysis of RT ascertained that different brain regions were recruited for domain-specific *vs*. domain-general cognitive control, and replicated this differentiation in the independent dataset of Humphreys and Lambon Ralph (2017). Below we elaborate on two issues relevant to our findings: (***i***) Semantic control as subpart of the broader semantic system; (***ii***) How do the gradient and topography approaches provide unique insights about cognitive control for semantic *vs*. non-semantic information?

### Semantic control regions as subpart of the broad semantic system

Decades of research have revealed that performing a semantic task engages widely distributed and functionally heterogeneous cortex (Binder *et al*. 2009; Martin 2015; Lambon Ralph *et al*. 2017). Based on a multitude of evidence, we proposed the *controlled semantic cognition* framework to account for the functional heterogeneity within the broad semantic system (Lambon Ralph *et al*. 2017). Under this framework, the ability to adaptively use semantic knowledge relies on two distinct neural machineries: (***i***) the brain regions that *represent* semantic knowledge *per se*, and (***ii***) the brain regions that *control* the retrieval of semantic information in a goal-directed fashion. Areas involved in *representing* semantics have been characterised as a ‘hub-and-spokes’ system, with the spokes being modality-specific cortex that encode embodied attributes (e.g., visual or motoric features) and the ventrolateral anterior temporal lobe (ATL) being a hub that encodes transmodal semantic meaning (Patterson *et al*. 2007; Lambon Ralph 2014). Over many years, there has been substantive evidence suggesting that transmodal concepts and modality-based concepts are underpinned by the ATL hub and modality-specific cortices (e.g., Pobric *et al*. 2010; Chiou and Lambon Ralph 2016, 2019).

Crucially, semantic control regions (e.g., the left IFG, pMTG, dmPFC) have fundamentally distinct functional roles from semantic representation regions. Damage to semantic control regions drives distinct behavioural deficits from damage to representation regions, causing two contrastive types of neurological disorder: semantic aphasia *vs*. semantic dementia (e.g., Jefferies *et al*. 2005; Jefferies *et al*. 2006; Jefferies and Lambon Ralph 2006; Jefferies, Hoffman, *et al*. 2008; Stampacchia *et al*. 2018; Thompson *et al*. 2022). Compared to the functional profile of semantic representation regions (e.g., the ATL), semantic control regions are more sensitive to varying task demands (Hoffman *et al*. 2010), more sensitive to the extent of difficulty in accessing meaning (Chiou *et al*. 2018; Gao *et al*. 2021), less affected by psycholinguistic variables (e.g., lexical familiarity/frequency/age of acquisition) and instead more affected by misleading contextual cues (Jefferies, Patterson, *et al*. 2008; Lanzoni *et al*. 2019). Thus, although co-activation of semantic control areas and semantic representation areas are frequently observed in the fMRI literature (particularly when a semantic/linguistic task is contrasted against a non-semantic task), there is a division between control regions and representation regions, as opposed to a homogenous semantic system.

### Cognitive control for different types of information through a topographical lens

When studying the neural basis of executive control, researchers usually create experimental contexts with minimal reliance on prior knowledge so that the task is maximally novel and challenging. Thus, stimuli conveying little semantics (e.g., scrambled shapes) and tasks probing non-linguistic processes (e.g., arithmetic) are generally favoured over lexical-semantic stimuli and semantic decisions. Because the key question is whether the MD network amplifies its reaction when confronted with a novel challenge, semantic knowledge is deemed ‘contamination’ that complicates the interpretation. However, semantic control is an intriguing borderline case that taps on the cognitive resources of *both* executive control and semantic meaning (i.e., goal-directed operation on semantics, requiring both selective attention and prior knowledge). Interestingly, our data revealed that semantic control took place physically in bridging cortical zones between the MD and default-mode systems, as revealed by the orderly topographical patterns observed in multiple sections of the brain. Crucially, our evidence of orderly arrangement lends further support to previous findings that the brain’s functional networks are spatially configured as highly reproducible topography and suggests the possibility that a brain region’s functional specialisation is sculpted by its neighbouring areas that supply input and receive output. Previously, Fedorenko *et al*. (2013) showed that the MD network has a reliable topography across individuals, lying next to modality-related regions. In a similar vein, the present results go a step further by demonstrating that semantic control regions emerge as the consequence of a general cortical motif – a brain region located in the intermediary strip between the MD system and default-mode system relays information between the two systems. We speculate that, as a result of its topographical loci, semantic control regions amalgamate the functional preferences of its neighbours, driving its specialised tuning of control for memory-based representations.

There has been mounting evidence that an area’s connectivity ‘fingerprint’ (i.e., all of its connections with rest of the brain) can be a key driving force that mechanistically shapes its functional profile (for discussion, see Mars *et al*. 2018). This connectivity-based framework has been used to explain how occipitotemporal sub-regions develop their preference for different visual entities, such as faces, animals, tools, places (e.g., Konkle and Caramazza 2017). This connectivity-based approach has also been employed to explore the MD system and its connectivity with auditory/visual-specific regions: Assem *et al*. (2021) and Tobyne *et al*. (2017) both showed that whether a subregion of the MD system had a bias towards auditory or visual stimuli hinged on its connectivity with auditory or visual regions. We speculate that connectivity may, in a similar vein, be the mechanism that drives the emergence of semantic control regions. Some previous resting-state connectivity evidence (Davey *et al*. 2016; Humphreys and Lambon Ralph 2017) and structural covariance evidence (Wang *et al*. 2018) have given support to our speculation: The two key sites of semantic control regions – the left IFG and pMTG – were found to be connected to both MD regions (e.g., the middle frontal gyrus and intraparietal sulcus) and default-mode regions (e.g., the ATL and the rostral subpart of angular gyrus).

More generally, our findings highlight the need to understand how topography/connectome shapes the functional relationships among regions/networks. As nicely delineated by Margulies *et al*. (2016), the principal gradient (from sensory-motoric to abstract-cognitive) is a powerful constraining force that shapes the positions of different conventional resting-state networks. The same orderly transitions are found across the brain, from sensorimotor areas, through MD areas, to default-mode areas. Although Margulies’s evidence is based on the topography of intrinsic connectivity at rest, it shows highly consistent patterns to task-based fMRI evidence in the present study (see also Wang et al., 2020). Together, these converging findings underscore the importance of understanding the functions of a brain region by situating it in the macroscale context of the whole-brain connectome.

## Acknowledgements

This research was funded by an MRC programme grant and intramural funding to MALR (MR/R023883/1; MC_UU_00005/18), a Sir Henry Wellcome Fellowship (201381/Z/16/Z) to RC, and a European Research Council Grant (771863 – FLEXSEM) to EJ.

## Footnotes

1 For completeness, we also identified areas tuned to the opposite interaction – greater response to visuospatial difficulty (Vis.-Hard > Vis.-Easy) *and* semantic easiness (Sem.-Easy > Sem.-Hard). Results are in Figure 4(A): This interaction identified various regions of the dorsal-attention system (the blue clusters: the bilateral frontal eye field and the superior/posterior parietal lobules), as well as many regions of the visual cortex.

2 The subdivision within the left IFG has been a matter of discussion (e.g., Fedorenko and Blank 2020). The IFG peaks we analysed here are the IFG’s anterior subpart (*pars orbitalis*) and its middle subpart (*pars triangularis*); however, a transition indeed existed in the IFG along its posterior to anterior axis, from visuospatial control, through semantic control, to visuospatial easiness (see *Supplemental Results 1*).

3 Although the default-mode network generally tends to be more active for cognitively less effortful contexts, it is an oversimplification to define this network as a ‘task-negative’ system. While in the present study we found that the default network became less active when confronted with semantic difficulty and visuospatial difficulty, it has been shown that this network has greater activation for behaviourally more difficult processes in perceptually-decoupled contexts (Murphy *et al*. 2018; Murphy *et al*. 2019). It has been suggested that the functional goal of default network is to sustain introspective processes, inducing episodic memory, semantic knowledge, and conscious thoughts (Smallwood, Turnbull, *et al*. 2021).

## Notes

**Conflict of interest:** The authors declare no competing financial interests.

### Competing Interest Statement

The authors have declared no competing interest.

### Summary of Updates

Revised on 16th August 2022, based on the reviewers' comments.

