## Supplemental Information for "A middle ground where executive control meets semantics: The neural substrates of semantic-control are topographically sandwiched between the multiple-demand and default-mode systems"

**Supplemental Stimuli**

The Semantic task included two conditions (Easy and Hard). A quadruplet of words were shown in each trial, containing an oddball (target) that was semantically inconsisent with the remaining three (Foil 1 – 3). Note, the target's location vaired randomly from trial to trial, equally likley to be in any of the four positions and reacted to by any of the four designated buttons in the actual experiment.

| Semantic Easy | | | | Semantic Hard | | | |
|---|---|---|---|---|---|---|---|
| Target | Foil-1 | Foil-2 | Foil-3 | Target | Foil-1 | Foil-2 | Foil-3 |
| Apple | Snow | Rain | Storm | Sprite | Gin | Vodka | Cider |
| Giraffe | Parcel | Packet | Package | Thistle | Pine | Maple | Birch |
| Ceiling | Lips | Mouth | Tongue | Delight | Anxiety | Depression | Delusion |
| Telephone | Week | Month | Year | Dog | Cat | Lion | Leopard |
| Bulldozer | Hand | Finger | Arm | Cosmos | Venus | Jupiter | Neptune |
| Antenna | Foot | Toe | Leg | Notepad | Pamphlet | Leaflet | Brochure |
| Elephant | Toilet | Loo | Lavatory | Pond | Brook | Stream | River |
| Butterfly | Street | Road | Avenue | Mick Jagger | John Lennon | George Harrison | Ringo Starr |
| Napkin | One | Two | Three | Trumpet | Guitar | Violin | Harp |
| Office | Mother | Father | Parent | Footprint | Trail | Path | Route |
| Tornado | Mug | Cup | Jug | Cello | Oboe | Piccolo | Tuba |
| Camel | Red | Blue | Green | Google | Facebook | Instagram | Twitter |
| Ape | Coffee | Tea | Water | Chicago | Berlin | Paris | Madrid |
| House | Monday | Tuesday | Wednesday | Philosophy | Zoology | Botany | Ecology |
| Moth | Dirt | Mud | Dust | Donald Trump | Gordon Brown | Tony Blair | David Cameron |
| Eyebrow | Sun | Moon | Star | Cabin | Castle | Mansion | Palace |
| Cyclone | Myself | Yourself | Himself | Smirnoff | Corona | Heineken | Guinness |
| Ladybird | Fork | Knife | Spoon | Prada | Reebok | Adidas | Nike |
| Galaxy | First | Second | Third | Belfast | Leeds | Reading | Edinburgh |
| Roof | Mine | Yours | Hers | Cristiano Ronaldo | Andy Murray | Novak Djokovic | Rafael Nadal |
| Sponge | Child | Toddler | Kid | Bedsheet | Blanket | Duvet | Quilt |
| Goose | Million | Billion | Trillion | Adele | Geri | Emma | Victoria |
| Daffodil | Shore | Beach | Coast | Boeing | Volvo | Honda | Audi |
| Nurse | Norway | Finland | Sweden | WhatsApp | Firefox | Safari | Chrome |
| Flame | Chin | Cheek | Jaw | Vodafone | NatWest | HSBC | Barclays |
| Owl | Bus | Truck | Train | History | Algebra | Calculus | Geometry |
| Fuel | Sheep | Horse | Cow | Neon | Silver | Nickel | Mercury |
| Lizard | Piano | Flute | Drum | Shield | Arrow | Dagger | Spear |
| Kitten | May | June | July | Seahorse | Dolphin | Walrus | Manatee |
| Shrimp | Coke | Pepsi | Fanta | Schoolyard | Laboratory | Gymnasium | Auditorium |
| Rocket | Ant | Bee | Wasp | Toenail | Pelvis | Ribcage | Skull |
| Puppy | London | Manchester | Bristol | Boston | Texas | Florida | Oregon |
| Dove | Pen | Paper | Rubber | Orion | Taurus | Virgo | Pisces |
| Bottle | Ear | Nose | Eye | Spider | Cricket | Mosquito | Fireflies |
| Otter | Teacher | Pupil | Student | Testicle | Ovary | Uterus | Cervix |
| Octopus | Actor | Athlete | Author | Dundee | Leicester | Exeter | Plymouth |

### Supplemental Results 1

In multiple places of the brain, the clusters of semantic control (red: Sem.-Hard > Sem.-Easy) were adjoined by visuospatial difficulty (blue cluster: Vis.-Hard > Vis.-Easy) and visuospatial easiness (green cluster: Vis.-Easy > Vis.-Hard), forming a consistent topography. Apart from the examples reported in the main article, we also found this topography in the left inferior frontal gyrus (IFG) and in the left lateral temporal cortex. To highlight the topography and avoid cluttered visual layout, anatomical masks (WFU_Pickatlas toolbox: Maldjian, Laurienti, Kraft, & Burdette, 2003) were used to confine visualisation of group-level results within the regions of interest.

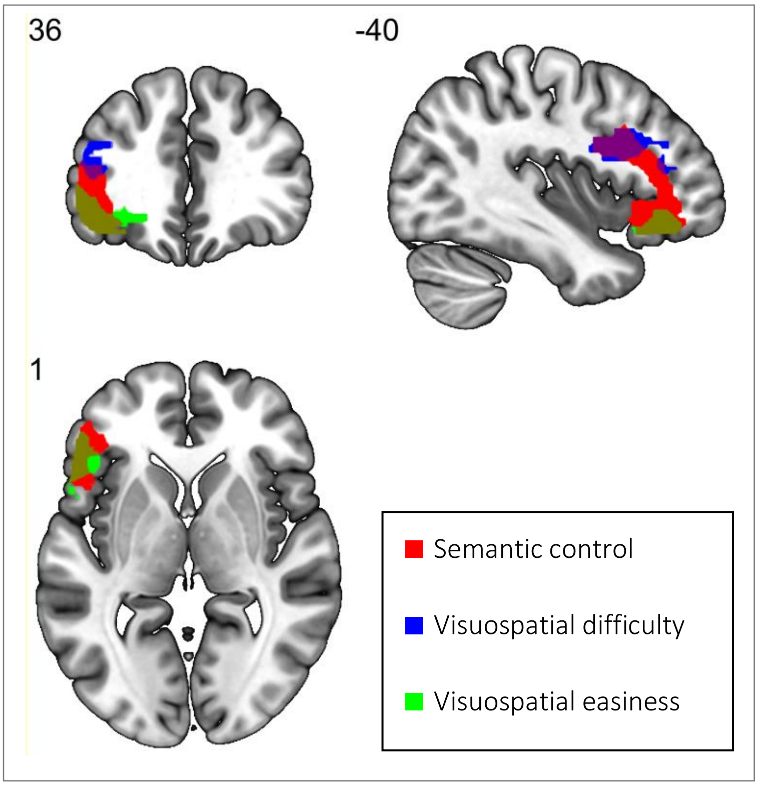

**(Supplemental Figure S1-1)**

Within the left IFG, the cluster of visuospatial difficulty was situated in the posterior subpart (*pars opercularis*), and visuospatial easiness was situated in the anterior subpart (*pars orbitalis*). Semantic control involved nearly the entire chunk of the left IFG and partially overlapped with the clusters of visuospatial difficulty and easiness. The middle section of the left IFG (*pars triangularis*) was almost exclusive to semantic control. Results were thresholded at $p < 0.001$ (voxel-wise) and $p < 0.05$ (whole-brain family-wise correction for clusters). Visualisation here was confined to only showing activation in the IFG mask.

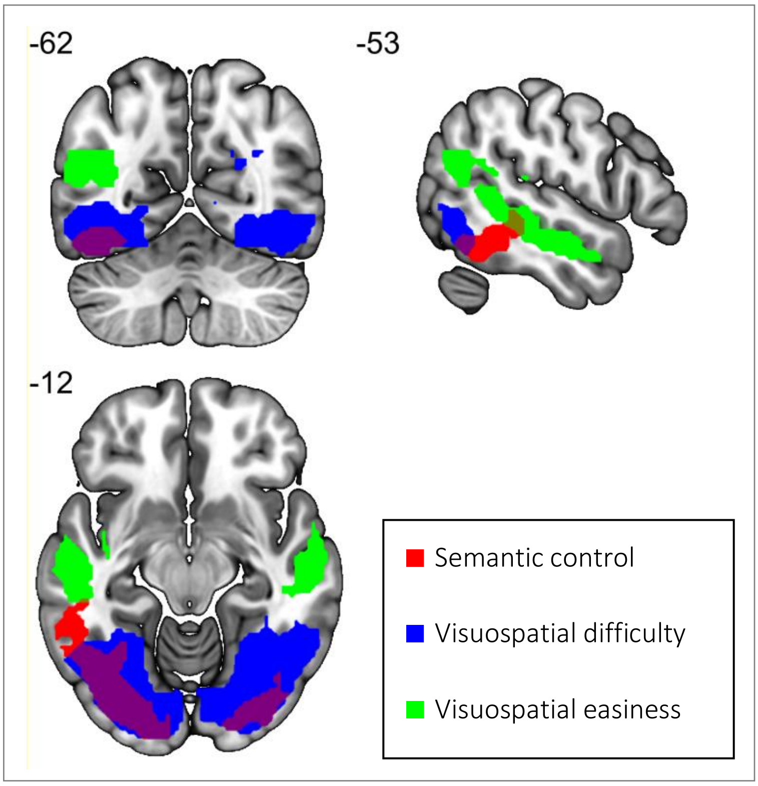

**(Supplemental Figure S1-2)**

In the bilateral occipital and temporal lobes, visuospatial difficulty involved primarily the ventral occipital cortices and inferior temporal gyri, while visuospatial easiness involved the superior temporal gyri. The Semantic control cluster was exclusive to the left hemisphere and situated in the posterior middle temporal gyrus (pMTG). Similar to other places of cortex, visuospatial difficulty and easiness flanked semantic control in the intermediate locus. Results were thresholded at $p < 0.001$ (voxel-wise) and $p < 0.05$ (corrected for family-wise error at the cluster level). Visualisation here was confined to only showing the occipital and temporal cortex.

**Supplemental Results 2**

In this analysis, we separately examined how 10 subregions of the multiple-demand (MD) system responded to semantic difficulty (Semantic-Hard > Semantic-Easy) and visuospatial difficulty (Visuospatial-Hard > Visuospatial-Easy). The 10 MD regions were based on the cortical parcels from the study of Fedorenko *et al.* (2013); in their group-level data, various frontoparietal areas, collectively forming the MD system, showed robustly heightened activation for taxing conditions relative to easier conditions. The MD network was segregated using anatomical/atlas landmarks, resulting in 10 cortical parcels (Figure S2-A and S2-B): the left/right middle frontal gyrus (red), left/right frontal eye field (green), left/right anterior/middle section of the parietal lobe (yellow), left/right anterior cingulate cortex (magenta), and left/right insula (blue).

The analysis detected a significant interaction involving all three factors ($F_{(4, 96)} = 5.33$, $p = 0.001$) – Difficulty Type (semantic/visuospatial), Hemisphere (left/right), Subregion (5 in each hemisphere). Results of post-hoc comparisons are shown in Figure S2-A and S2-B. As the Figure illustrates, hemispherical difference modulated the MD system's reaction to the two types of difficulties. Simple effects showed that the semantic difficulty effect was greater in the left hemisphere (consistent with the left-lateralisation of language functions); the semantic effect was also greater in prefrontal areas (mid-frontal gyri, insula, ante-cingulate cortex) than in those regions typically implicated in saccade and visuomotor functions (frontal eye fields, parietal areas).We interpret this pattern (more prefrontal activation, less visuomotor regions) as reflecting the Semantic task's emphasis on cognitive control over visuomotor processes. Simple effects also showed that the visuospatial difficulty effect reached statistical significance in every subregion of the MD system and in both hemispheres. It is worth noting that comparing the sizes of semantic difficulty versus visuospatial difficulty is not appropriate, because of differential baselines in behavioural data (namely, the reaction times of Semantic-Easy were slower than those of Visuospatial-Easy).

Next, we situated the activation driven by semantic control (i.e., Semantic-Hard > Semantic-Easy) in generic contexts of various semantic processes (Figure S2-C) and executive-control processes (Figure S2-D). We used the function of term-based meta-analysis of NeuroSynth, searching for 'semantic' (based on 4,031 fMRI studies and 40,030 activation points) and 'executive' (based on 786 fMRI studies and 28,937 activation points). Results of the Meta-analysis clusters were thresholded at $q < 0.01$ (corrected for voxelwise false discovery rate). As shown in Figure S2-C and S2-D, the activities of semantic control (red clusters) are compared with the meta-analysis of semantic processes (yellow clusters in S2-C) and executive-control (blue clusters in S2-D).

In Figure S2-C, the semantic control clusters partially overlapped with the expansive set of areas involved in miscellaneous semantics-related processes. Note that the semantic control clusters (identified by the present study) overlapped with the meta-analysis in the left prefrontal cortex and posterior mid-temporal gyrus/pMTG (two established areas in the semantic control literature) while they did not overlap in large chunks of the anterior temporal cortex (ATL). This pattern is consistent with the literature that *semantic control* relies on the left prefrontal cortex and pMTG while *semantic representation* depends on the ATL. In Figure S2-D, the semantic control clusters also partially overlapped with the meta-analysis of executive process; the most obvious difference is the right posterior parietal lobe that semantic control did not engage this region, which has been well-stablished in the literature of spatial attention.

Taken together, we found that activation was heightened in most subregions of the MD system when tasks became difficult, observed both for the Semantic or Visuospatial task. It is, however, important to note that the MD network's role in a language task has been demonstrated to reflect maintaining attentional focus on the task, rather than semantic processes *per se* (Diachek, Blank, Siegelman, Affourtit, & Fedorenko, 2020). Moreover, semantic control was relativley lateralised to MD areas in the left hemisphere, and relied more on prefrontal areas (relative to parietal areas), while visuospatial control robustly activated every subregion. Collectively, the data are consisent with MD regions' prominent role in non-language tasks and their ancillary role in language tasks.

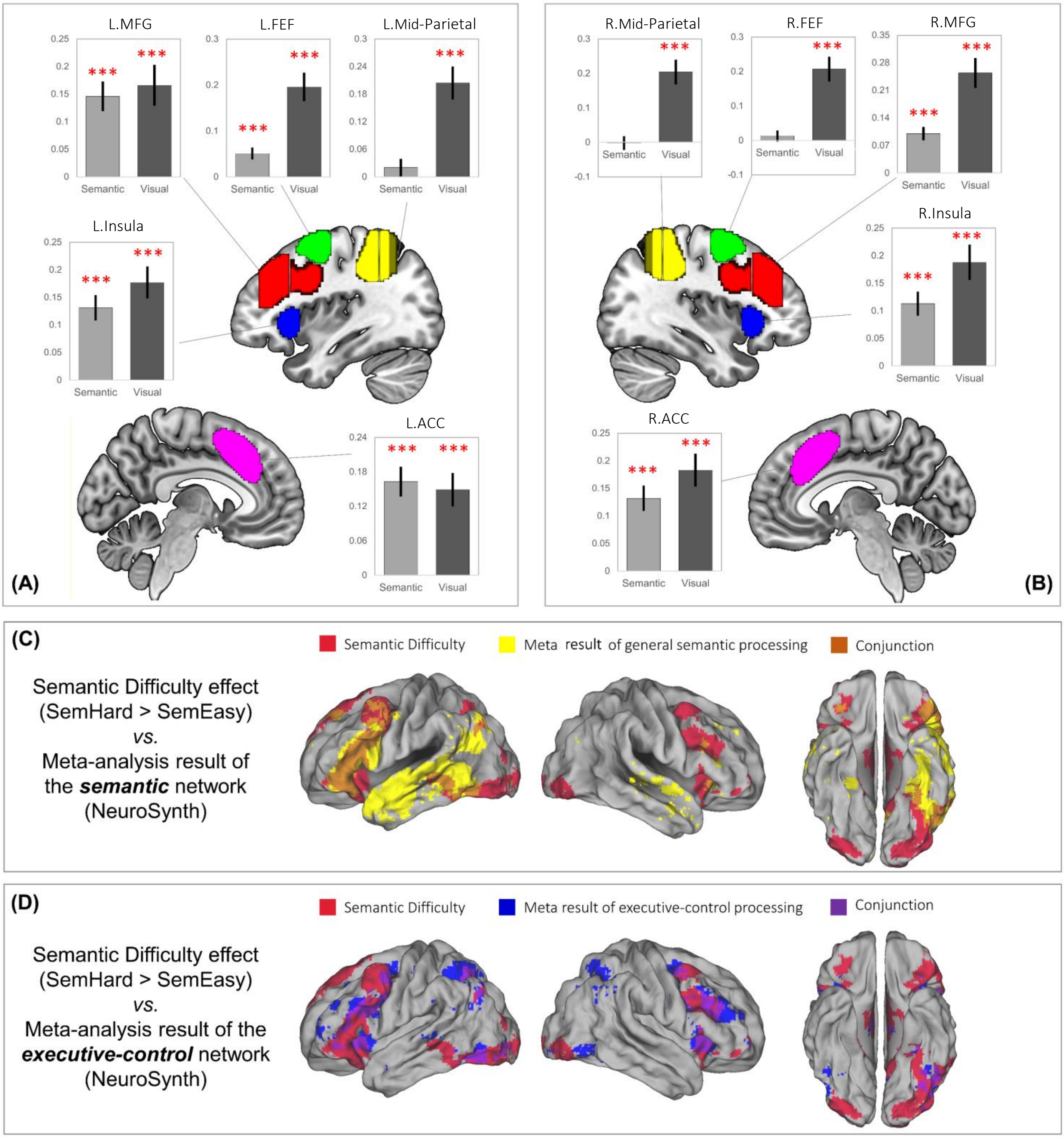

**(Supplemental Figure S2)**

**Supplemental Results 3**

We contrasted the task-induced activation of Semantic-Hard *vs.* Visuospatial-Hard, given that the two conditions had statistically matched performance levels. Results of whole-brain analysis was thresholded at $p < 0.001$ (voxelwise intensity) and $p < 0.05$ (FWE-corrected for cluster-level multiple comparisons). As shown in Figure S2, the contrast 'Semantic-Hard > Visuospatial-Hard' identified a set of regions – the left inferior frontal gyrus (IFG), left superior temporal gyrus, bilateral temporal ATLs (temporal poles), bilateral supramarginal gyri, medial prefrontal cortex, medial temporal lobes (including the hippocampus). Collectively, these regions exhibited the typical topography of the language network (e.g., Diachek et al., 2020); their topography also resembled the typical patterns of transmodal areas that are situated at the 'abstract-cognitive' end of principal cortical gradient (Margulies *et al.*, 2016). The reverse contrast 'Visuospatial-Hard > Semantic-Hard' revealed brain areas well-documented in the literature of spatial attention, saccade, and various externally-directed visuomotor processes – the superior parietal lobule, intraparietal sulcus, frontal eye field, as well as visual cortex. The topography of these regions resembled the multiple-demand network (Fedorenko, Duncan, & Kanwisher, 2013), as well as the dorsal-attention network and visual network (Yeo et al., 2011).

It is worth emphasising the functional divergence (semantic control *vs.* semantic representation) among regions involved in language/semantic processing (for review, see Lambon Ralph, Jefferies, Patterson, & Rogers, 2017). To detect such difference, different contrasts had to be used: ***First***, the comparison of 'Semantic-Hard > Visuospatial-Hard' revealed the typical configuration of the entire semantic network, which included both semantic representation regions (e.g., the ATLs) and semantic-control regions (the IFG and pMTG). Representations and control regions both preferred semantics to visuospatial processing, hence they were both activated by this contrast. ***Second***, to identify areas specifically tuned to semantic control, a contrast that differentiated the hard level of semantic operation from the easy one would be necessary (namely, Semantic-Hard > Semantic-Easy); this contrast selectively identified the subsystem of semantic-control (the IFG and pMTG) while sparing the subsystem of semantic representation (the ATL).

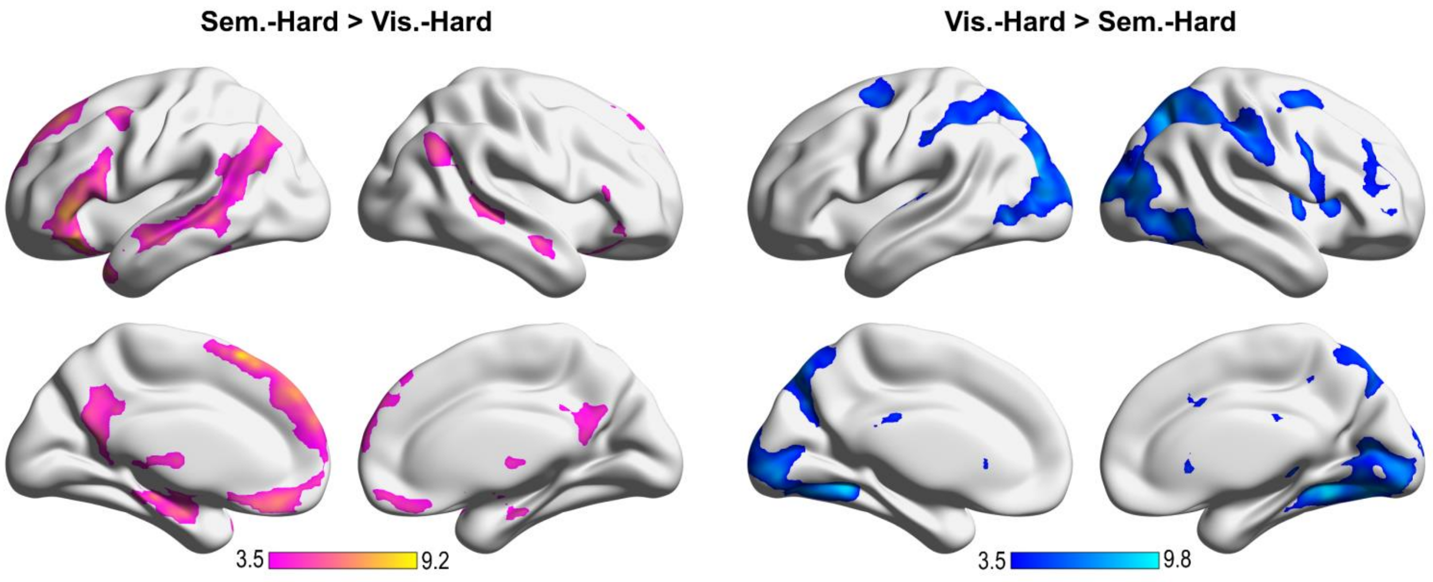

**(Supplemental Figure S3)**
